# Using image-based haplotype alignments to map global adaptation of SARS-CoV-2

**DOI:** 10.1101/2021.01.13.426571

**Authors:** Tom W. Ouellette, Jim Shaw, Philip Awadalla

## Abstract

Quantifying evolutionary change among viral genomes is an important clinical device to track critical adaptations geographically and temporally. We built image-based haplotype-guided evolutionary inference (ImHapE) to quantify adaptations in expanding populations of non-recombining SARS-CoV-2 genomes. By combining classic population genetic summaries with image-based deep learning methods, we show that different rates of positive selection are driving evolutionary fitness and dispersal of SARS-CoV-2 globally. A 1.35-fold increase in evolutionary fitness is observed within the UK, associated with expansion of both the B.1.177 and B.1.1.7 SARS-CoV-2 lineages.

## Main

Identifying adaptive processes associated with phenotypic change is critical to understanding how organisms evolve over time and identifying targets for clinical therapy. During the SARS-CoV-2 pandemic, a large number of complete RNA genome sequences from a number of populations have now been captured^1^, and we are arguably well-powered to determine if the SARS-CoV-2 genome is subject to selection globally, or within various regions, and changing over time. Identifying populations of SARS-CoV-2 genomes or haplotypes subject to positive selection over time has implications for monoclonal antibody or vaccine development and for understanding phenotypic changes, such as altered virulence or immunogenicity over time^2,3^

Many population genetic and phylogenetic methods exist to discriminate among multiple evolutionary processes or models, but the majority of methods utilize summary statistics that reduce sequence information into a condensed numerical or graphical output^4,5^. Traditionally, these methods aim to identify the distribution of nucleotide variation expected under neutrally evolving scenarios, often assuming or requiring free recombination and constant population sizes, and make interpretations following the detection of outliers^6^. In contrast, phylogenetic approaches rely on recombination-free models to make inferences, but are dependent on having numerous recurrent events within a given time interval such that excesses beyond neutral expectations are captured, often limiting their application to highly mutating genomes or very old evolutionary histories^6^.

Deep learning methods provide an opportunity to exploit the information stored in haplotype alignments by framing evolutionary analysis as an image recognition problem^7–9^. Given the multi-dimensional and complex nature of certain evolutionary processes, developing statistical and probabilistic models with reasonable parameters is often intractable. Deep learning provides an opportunity to encode the complexity of evolution in tractable simulation-driven prediction tools that do not require explicit parameterization^7,9^. Early applications of deep learning in population genetics have generated accurate predictions of introgression, recombination rate, and selection, but have largely focused on recombining, germ-line data from human populations or single locus analyses^7,9–11^.

Here, we developed a convolutional neural network (CNN) combined with a recurrent neural network (RNN) approach (see Online Methods), called image-based haplotype guided evolutionary inference (ImHapE), to detect and quantify selection in non-recombining expanding populations with available genomes, such as SARS-CoV-2 (Figure 1a). In summary, ImHapE involves four main steps, (i) simulating forwards-in-time processes in small genomic windows of length *N*, (ii) training a CNN on aligned genomic windows of length *N*, (iii) simulating complete 29903 bp genomes and performing sliding window analysis using trained CNN and other population variant allele frequency summaries such as Tajima’s *D* (TajD)^12^ and Fay and Wu’s *H* (FWuH)^13^, and (iv) training a long short-term memory (LSTM) recurrent neural network (RNN) to classify positive selection and quantify evolutionary fitness (1 + s) in population of non-recombining genomes at a given time point (Figure 1b). Although a sliding window step seems unusual for applications within a non-recombining context, we chose to integrate a sliding window step as simulation and training a CNN on full genome alignments, e.g. 29903 bp such as SARS-CoV-2, is computationally expensive. By performing sliding window analysis first and then making inferences using an LSTM, we reduce the sequence length while still capturing the relative differences in fitness across populations subject to different strengths of positive selection. In addition, we integrate existing population genetic statistics, such as TajD and FWuH, at the sliding window step to improve model performance by combining multiple features with our CNN estimates. Our general framework can be adapted and applied to any non-recombining population where aligned haplotype information is available such as somatic tissues or cancers.

**Figure 1.**
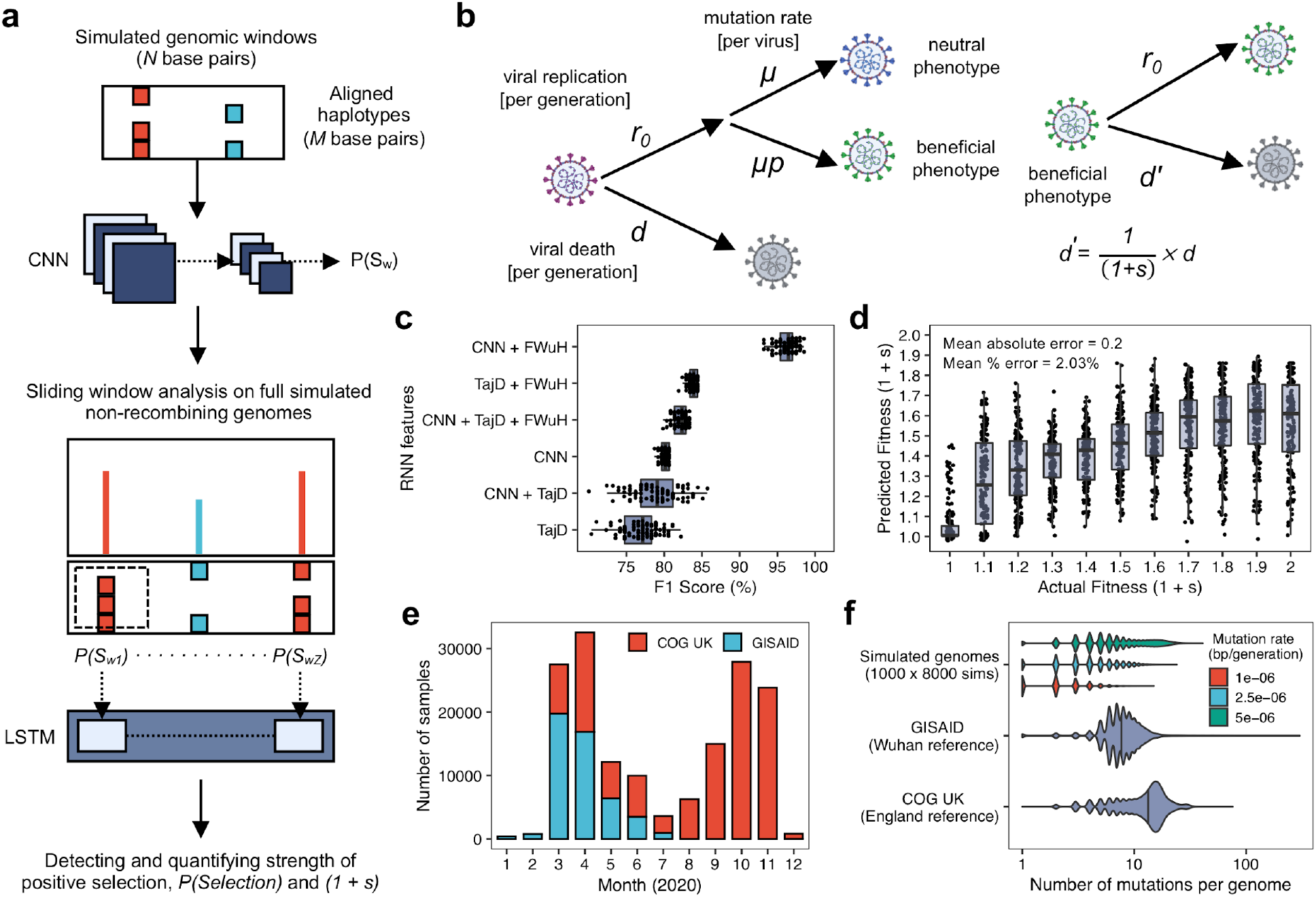
Inferring selection in non-recombining viral genomes through image-based haplotype alignments. **a**, Positive selection can be identified through a combined simulation-based convolutional neural network (CNN) and recurrent neural network (RNN) approach. Because training on full genome alignments is computationally expensive, information from full genomes can be accounted for by performing sliding window analyses using CNN estimates, *P(Sw)*, and analyzing the sliding window estimates with recurrent neural networks (RNN) such as a long short-term memory (LSTM) network. In addition, common population genetic statistics, such as Tajima’s *D*^12^ (TajD) and Fay and Wu’s *H*^13^ (FWuH) can be integrated at the sliding window stage to improve inferences. **b**, To generate training data for SARS-CoV-2 evolutionary inference, we simulated exponentially growing populations of one hundred thousand non-recombining viruses. The simulated viral populations grew exponentially with a replication rate (r0) and death rate (d). Mutations were Poisson distributed with neutral mutations arising at a rate μ and beneficial mutations arose at a rate μp where p was the probability of a mutation being beneficial. Viruses with a beneficial mutation had a reduced death rate (d’) that was inversely scaled by the fitness of the mutation (1 + s). All parameters used for simulations, and a complete simulation overview, are provided in Online Methods **c**, Positive selection classification performance across individual and combined sliding window features. The F1 score (harmonic mean of precision and recall) was computed 100 times across subsamples of 100 neutral (n = 3000) and positive (n = 3000) simulated genomes. Combining CNN and FWuH sliding window estimates resulted in the strongest performance. **d**,. Actual fitness versus predicted fitness using an ensemble of LSTMs trained using different combinations of CNN, TajD, and FWuH features. **e**, Distribution of GISAID samples (March to July 2020) and COG UK samples (April to December 2020) retrieved for this study. **f**, Comparison of mutations per genome in simulated genomes (1000 genomes x 8000 simulations), GISAID samples (mutations called using Wuhan reference genome), and COG UK samples (mutations called using an England reference genome).

To first train our CNN to detect positive selection in genomic windows of length *N*, we simulated exponentially growing viral populations, with a final population size of one hundred thousand, evolving under a random-birth death process (Figure 1b; Online Methods). Both neutrally evolving populations and populations subject to selection grew under exponential growth with positively selected subpopulations growing at a faster rate dependent on fitness (Figure 1b; Online Methods). We defined increased fitness as a reduction in death rate such that a fitness (1 + s) of 2 was equivalent to a 50% reduction in the viral death rate in the beneficial virus population (Figure 1b; Online Methods). For each simulation, we sampled alignments of 200 genomes from the simulated viral population. Using 8000 alignments from both neutral evolution and positive selection simulations, we trained a binary CNN to return a classification score or probability estimate for positive selection in a given genomic window of length *N* (see Methods). We evaluated multiple genomic, or sliding, window sizes (see Methods) and found a genomic window length of 2500 bp to be sufficient to ensure separability of site frequency distributions from simulated populations subject to neutral or positive selection (Supplementary Figure 1). We achieved 95% recall and 99% precision for predicting positive selection in genomic windows of 2500 bp (Supplementary Figure 2). Finally, to make inferences in full 29903 bp genomes, we performed sliding window analyses across approximately 10,000 simulated genomes and then trained a long short-term memory (LSTM) RNN to predict if the genome was under positive selection and the population fitness (Online Methods). Half of the genomes were simulated as neutral and the other half were subject to positive selection with fitnesses ranging from 1.1 to 2. Evaluating all combinations of sliding window features (CNN estimates, TajD, and FWuH; Online Methods), we found that individually the CNN sliding window estimates outperformed, but combining both CNN and FWuH resulted in the highest F1 score of 95% (Figure 1c; Supplementary Figure 3). The combined CNN and FWuH model was used for classification of positive selection in empirical data. In addition to classifying positive selection, we built a model to quantify the evolutionary fitness in empirical data. To optimize quantitative estimates, we combined the top performing RNN models into an ensemble using a gradient-boosted machine and achieved a mean percentage error of 2% and a mean absolute error of 0.2 (Figure 1d; Supplementary Figure 3). We note that we do slightly overestimate lower fitnesses (1.1 − 1.5) and slightly underestimate higher fitnesses (1.5 − 2) but maintain reasonable fitness estimates of 1 for the majority of neutral simulated genomes (Figure 1d).

We applied our trained and validated CNN/RNN model to empirical global SARS-CoV-2 data from March to July 2020 (GISAID^1^) and to United Kingdom data from April to December 2020 (COG UK^14^). To ensure accurate inferences, we applied a conservative filtering step removing any samples with excessive genome masking relative to the overall sample distribution (Supplementary Figure 4; see Online Methods). After filtering, we retained 28,350 SARS-CoV-2 genome sequences from GISAID and 66,495 from COG UK for analysis (Figure 1e). We called mutations in the GISAID data using the Wuhan reference genome (NCBI RefSeq: NC_045512) to examine positive selection during early expansion of SARS-CoV-2 and called mutations in the COG UK data using a an England reference genome (COG UK: England/LOND-D51C5/2020) from April to study evolution post-fixation of the D614G haplotype^15^. As well, the mutation rate of SARS-CoV-2 is estimated to be 22.8 mutations per genome per year^16^ and we ensured our mutation rate was calibrated so that simulated genomes had similar numbers of mutations per haplotype alignment as empirical data (Figure 1f).

By pooling SARS-CoV-2 samples across specific geographic regions and time points for analysis, we find evidence for positive selection in all populations but with varying evolutionary fitnesses (fitness ranging from 1.05 to 1.5) in Asia, Europe, and North America from March to July 2020 (Figure 2a). The evidence for positive selection is observed despite the fact that all of our models incorporate population growth using empirically observed growth parameters (Online Methods). In European and North American samples, we observe an elevated but reduced fitness in July relative to earlier time points (Europe: March = 1.42, July = 1.05; North America: March = 1.40, July = 1.27), that occurred following fixation of the ubiquitous D614G haplotype^15^. Although fitness decreased over time in the continental data, fitness was still estimated larger than neutral (> 1) even under an exponential growth model.

**Figure 2.**
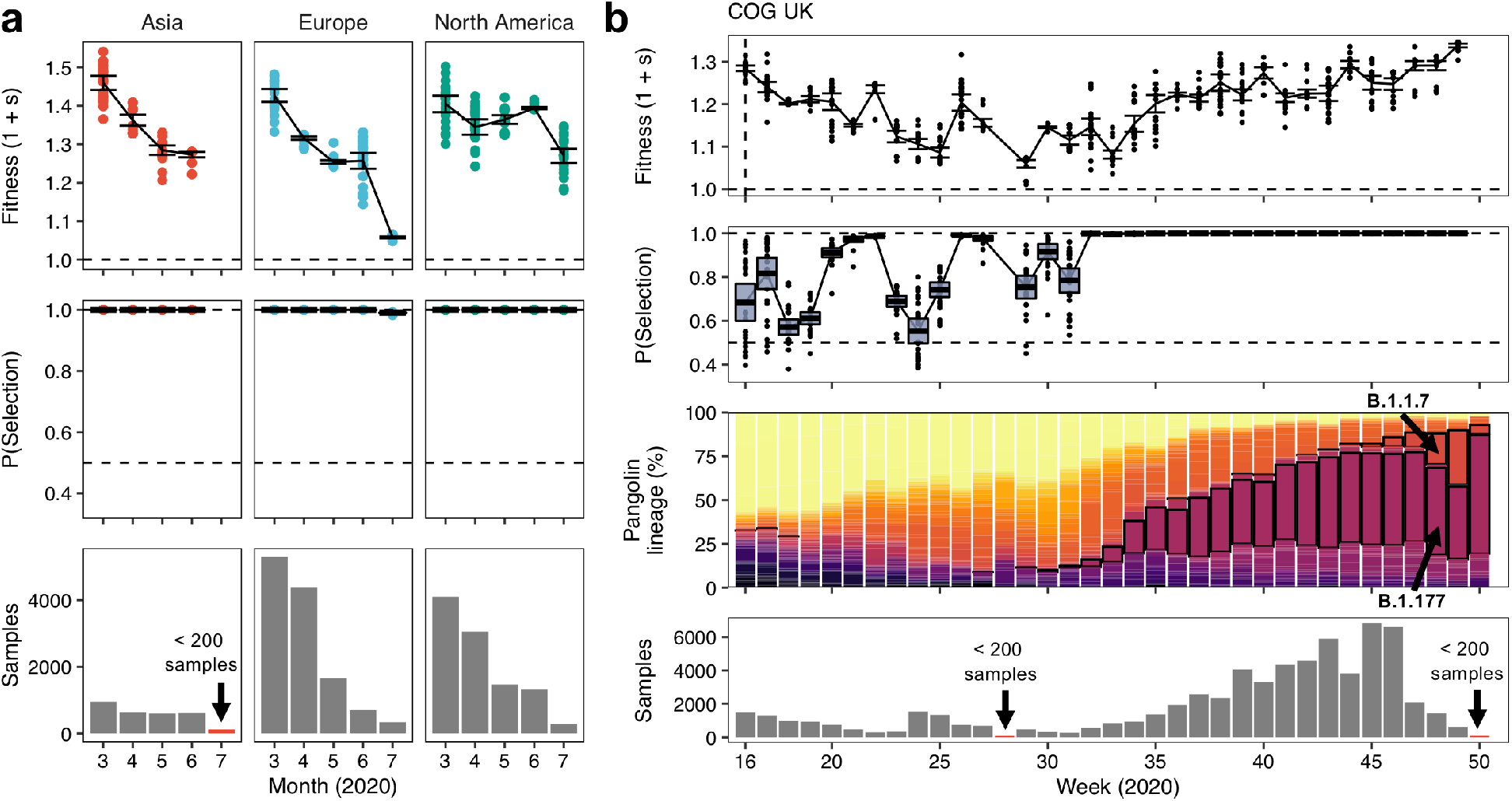
Detecting and quantifying positive selection globally and in the United Kingdom SARS-CoV-2 population using a combined CNN and LSTM based approach. **a**, We analyzed positive selection in 28,350 samples from GISAID^1^ from March to July 2020 in Asia, Europe, and North America. For each month with at least 200 samples (CNN alignment size), we computed both the fitness (1 + s) and probability of positive selection, P(Selection), across 25 subsamples of 200 samples from the empirical data. The upper and lower bands for both fitness and P(Selection) represent the upper and lower 95% confidence interval across the 25 subsamples. **b**, To analyze selection following the emergence of the D614G haplotype in early 2020, we called mutations using a reference genome from April 2020 (COG UK identifier: England-LOND-D51C5-2020) rather than the Wuhan reference genome. The fitness following week 29 was correlated with an expansion in the B.1.177 lineage (pangolin lineage) within the United Kingdom (COG UK data). In addition, the new emerging variant B.1.1.7 showed expansion in conjunction with a fitness increase following week 46.

The COG UK data provided a more thorough application of our model as abundant SARS-CoV-2 sequence data was available at a weekly time-scale (Figure 2b). We observe large variability in probabilities of positive selection at early time points but observe a consistent increase in fitness from week 29 (fitness = 1.05) to week 49 (fitness = 1.34).The fitness increase following week 29 was coincident with an expansion in the B.1.177 lineage (proportion at week 29 = 0%; week 49 = 41%) within the United Kingdom (COG UK data) and, in addition, the expansion of the new emerging variant B.1.1.7 following week 46 (proportion at week 46 = 5%; week 49 = 31%). The B.1.177 lineage characterized by an A222V spike protein mutation has been expanding across multiple European countries since its emergence in early summer of 2020^17^. Although there has been uncertainty around the phenotypic advantage of the B.1.177 lineage, our fitness estimates coincident with B.1.177 expansion suggest this lineage, and the corresponding A222V mutation, may provide an increased transmissibility. With that said, the emergence of the new B.1.1.7 lineage, characterized by 8 novel mutations (2 deletions and 6 non-synonymous)^18^, and corresponding increase in fitness following week 46 likely suggests the B.1.1.7 lineage may be fitter relative to the existing lineages circulating within the United Kingdom.

Both the continental and the COG UK data profiling is an excellent example of how the simulation-based CNN/RNN tools can track selective differences among viral clones almost in real time. Inferring selective differences contributes to the understanding of potentially clinically impactful pathogens that can change the virulence and infection rates globally. With as few as 200 samples within a region or a time-point, we can see significant shifts in estimates of evolutionary fitness. We acknowledge that inferences can be confounded by multiple limitations such as ascertainment or sampling bias due to differences in region-specific sequencing strategies and lockdown measures, or superspreader events that artificially augment specific lineage frequencies. With that said, we analyzed selection across pooled continental or country (UK) data to reduce biases driven by artificial over representation of lineages in specific locales. Overall, simulation-based CNN/RNN approaches provide a flexible framework for evolutionary inference, and we foresee further development and refinement of these approaches with application to other viruses and single-cell cancer data.

## Supplementary figures

**Supplementary Figure 1.**
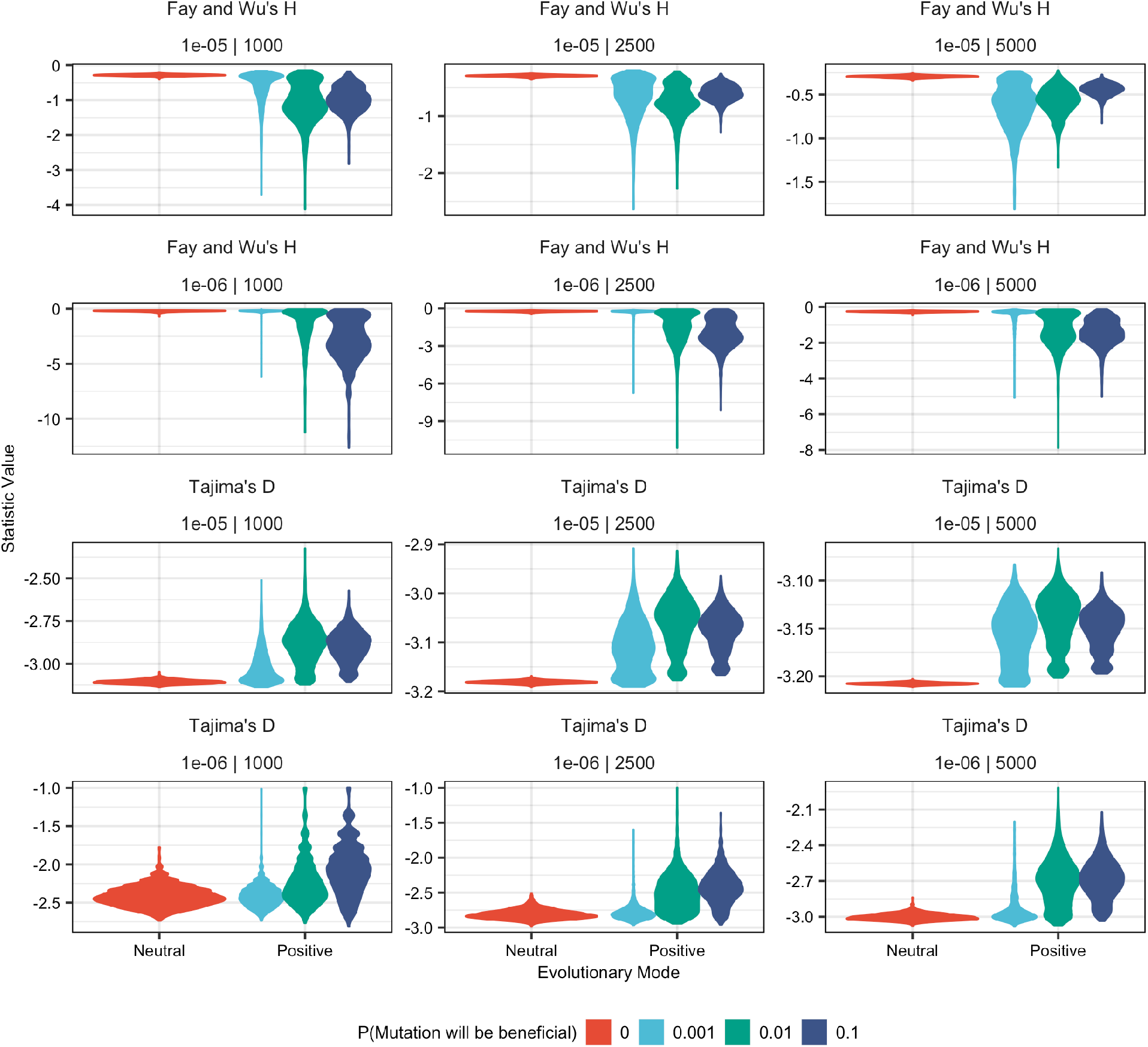
Evaluation of Tajima’s D (TajD) and Fay and Wu’s H (FWuH) across a range of window sizes (1000, 2500, 5000), mutation rates (1e-5, 1e-6 per base/gen), and probability of beneficial (0.001, 0.01, 0.1). To evaluate the separability of neutral and positive evolutionary models in simulated genomic windows, we simulated 2000 neutral and positive viral populations per parameter combination (window size, mutation rate, and probability of beneficial) and computed established site-frequency based summary statistics TajD and FWuH to determine which window size would give sufficient site frequency variation information.

**Supplementary Figure 2.**
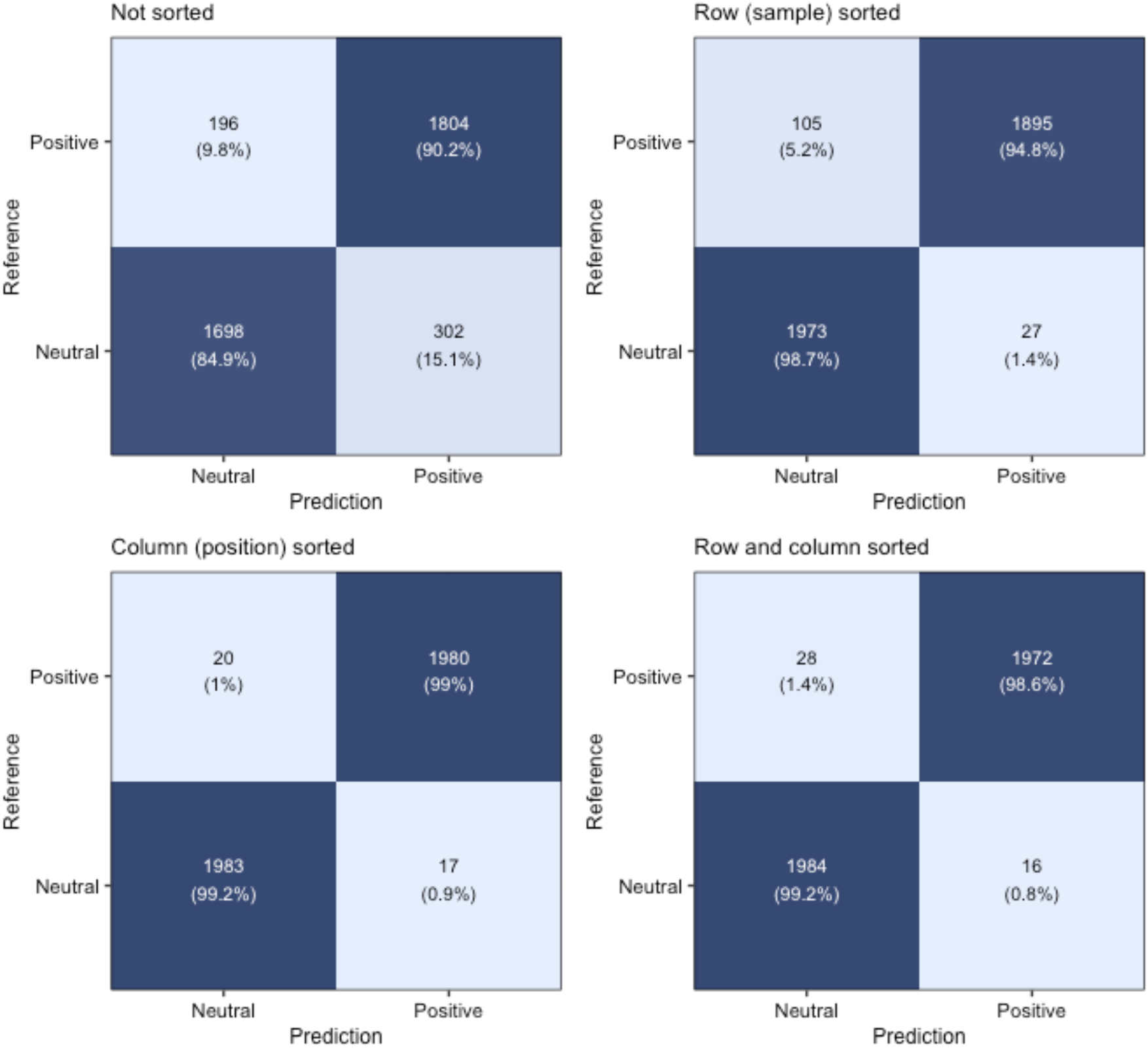
Performance of CNN predicting positive selection in genomic windows of 2500 base pairs. Previous studies in recombining organisms have shown that row (sample) and column (position) sorting of haplotype image-based alignments can improve inference via a CNN^7^. To evaluate the impact of sorting in the non-recombining context, we simulated 8000 neutral evolution and 8000 positive selection viral populations in genomic windows of 2500 bp per sorting method and trained a CNN to predict if positive selection was occurring. We found that both row and column sorting improved model performance, both precision and recall. With that said, we chose to utilize only row sorted models in our sliding window analysis as it retained the positional information (no column/positional sorting) while maintaining strong recall and precision.

**Supplementary Figure 3.**
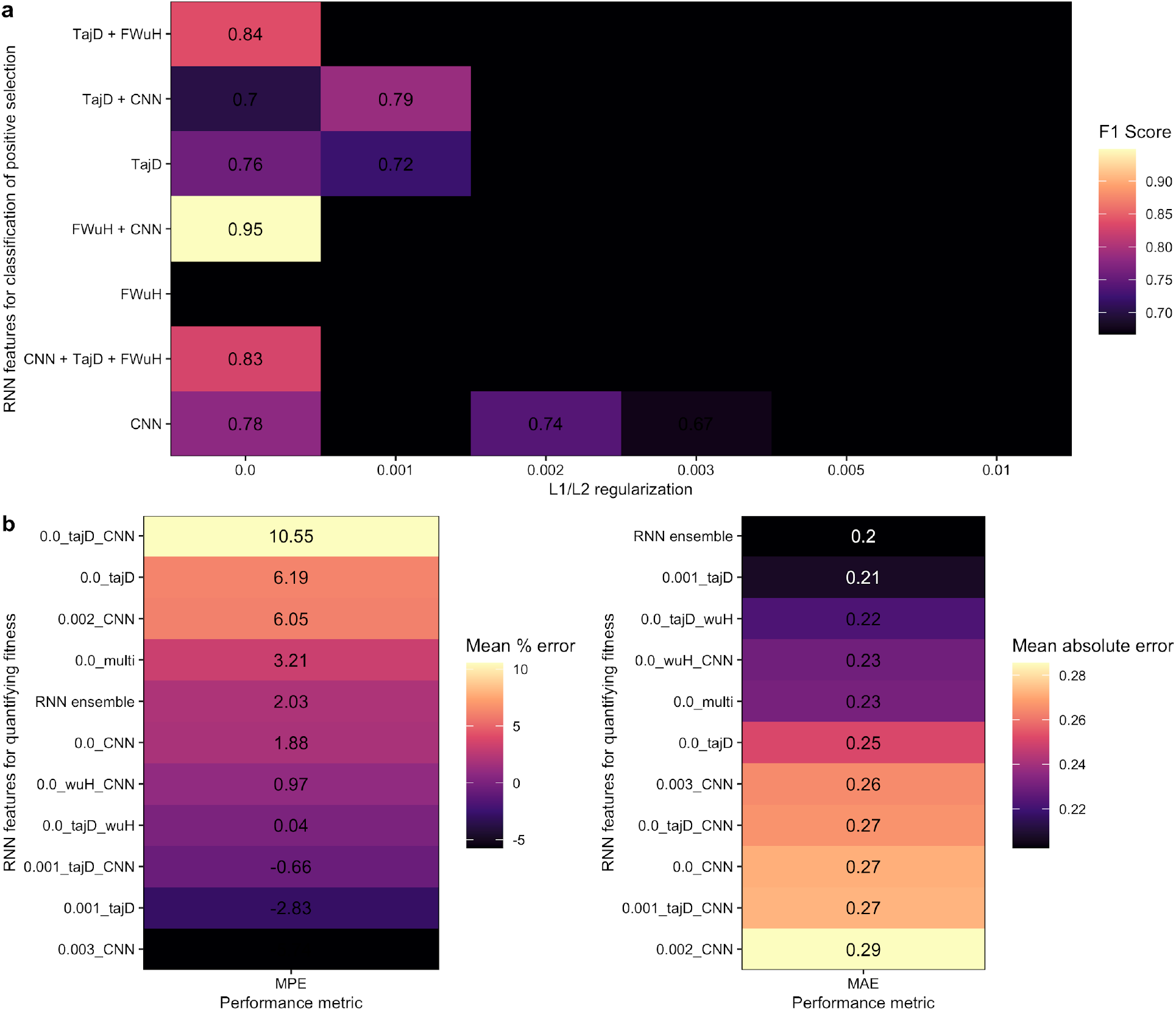
Selecting top performing models based on feature combinations and regularization of LSTM. To choose an LSTM-based model for predicting positive selection in empirical data, we first simulated 5000 neutral and 5000 positively selected viral populations and performed sliding window analysis using our CNN trained on 2500 bp genomic windows with row sorting. In addition, we computed TajD and FWuH at each sliding window. We evaluated all feature combinations (CNN, FWuH, and Tajima’s (TajD) across a range of L1/L2 regularization strengths. **a**, For classifying positive selection, we chose the top scoring model based on the F1 score (harmonic mean of precision and recall) in a holdout set of 3000 neutral and 3000 positive selection simulated populations. The F1 score was taken as the mean estimate across 100 subsamples of the data where positively selected samples were stratified by fitness. **b**, To choose an LSTM to quantify fitness, we computed both the mean percentage error and mean absolute error in a holdout set of 175 simulated genomes per fitness (fitness of 1 to 2, step size 0.1). We found that an ensemble of the top 10 models (all models evaluated here) outperformed all other models based on the mean absolute error while also maintaining a low mean percentage error.

**Supplementary Figure 4.**
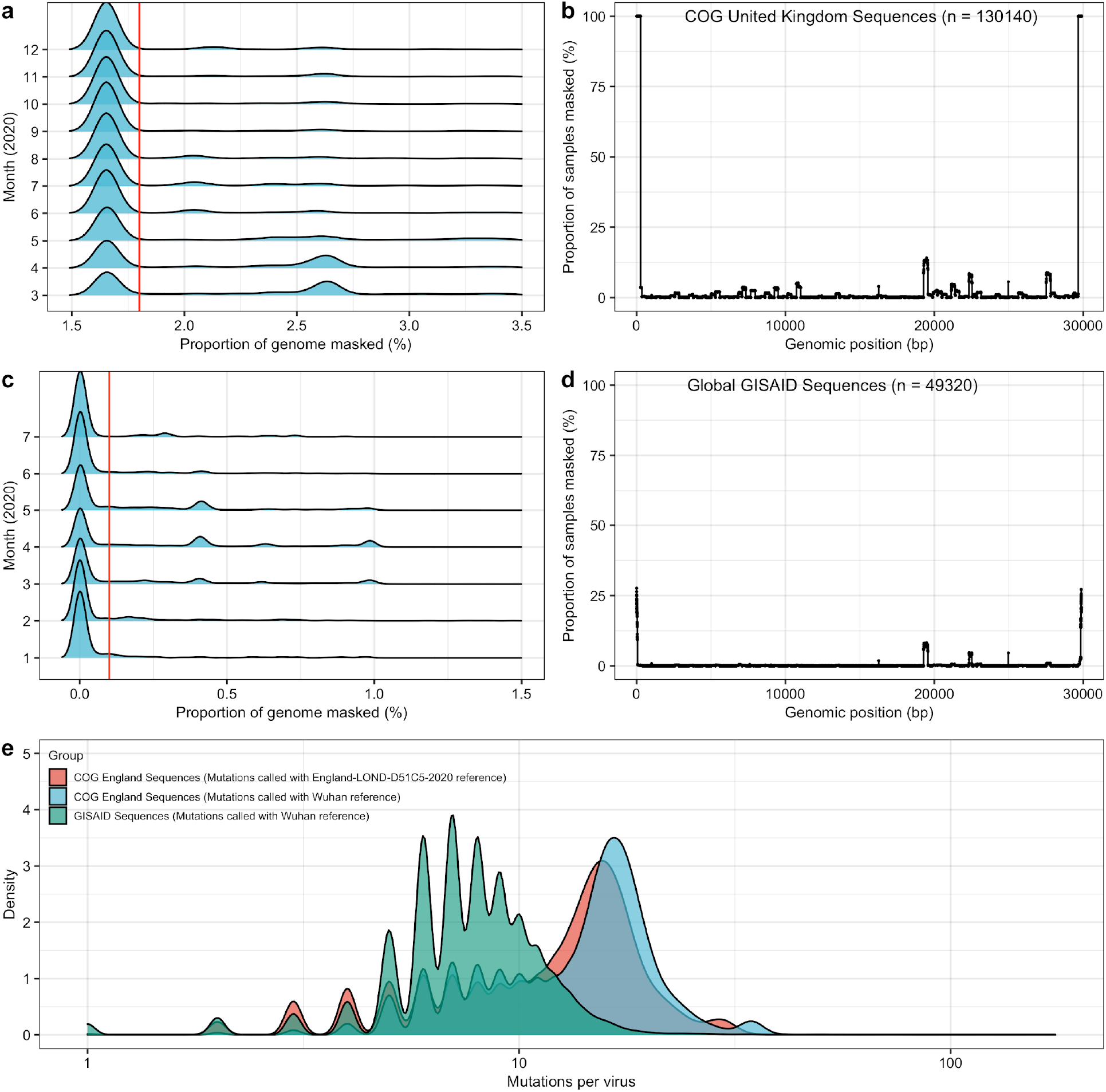
General quality control and sample filtering for GISAID and COG UK data. To ensure accurate inferences were being made in empirical data, we set a threshold for sample inclusion based on the proportion of genome masked. To set the threshold for the proportion of genome masked, we set simple cut-offs based on the entire sample distribution. For COG UK data, **a**, we only accepted samples with less than 1.8% of the genome masked, and **b**, did not find an excessive number of sites with masking in a genome-wide scan. For GISAID data, **c**, we only accepted samples with less than 0.1% of the genome masked, and **b**, did not find an excessive number of sites with masking in a genome-wide scan. **e**, We compared mutational counts in GISAID data (from January to August 2020) and COG UK data (from March to December 2020). In particular, we ensured that mutation counts using the LONG-D51C5 reference genome from April 2020 resulted in a smaller number of mutations on average then mutations called in COG UK data using the Wuhan reference from December 2019.

## Acknowledgements

We would like to thank GISAID and COG UK for aggregation, organization, and management of global SARS-CoV-2 sequences. We are particularly grateful to Jared Simpson, Marie-Julie Fave and Stephen Wright for comments and suggestions that improved analyses and the manuscript.

## Data/Code Availability

All SARS-CoV-2 sequences were retrieved from GISAID (https://www.gisaid.org/) and the COG UK consortium (https://www.cogconsortium.uk/). Code for this study is freely available under a GPL-3.0 license and can be found at https://github.com/tomouellette/ImHapE.

## Methods

### Modeling SARS-CoV-2 evolution

We modeled and simulated SARS-CoV-2 evolution as an exponentially growing, non-recombining population with a final population size of one hundred thousand. We applied our simulation framework to study SARS-CoV-2 under different evolutionary scenarios with increasing strengths of selection. The framework was modeled as follows. Mutations occurred randomly during each round of replication and were poisson distributed with neutral mutations occurring at a rate of *µ* and beneficial mutations occurring at a rate *µp*, where *p* is the probability a mutation was positively selected. Given the small genome size of SARS-CoV-2, we allowed for back mutations at neutral sites. Overall, the population evolved under a random birth-death process with a replication rate (*r*) and a death rate (*d*). To incorporate positive selection, viruses with beneficial mutations had increased fitness (*1 + s*) that inversely scaled the death rate. Therefore, viral populations grew exponentially based on the following function (where *N*_*t*_ is the population size at time *t*).

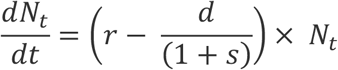

We chose to inversely scale the death rate with fitness as many mildly pathogenic viruses are known to become less virulent over time and have longer survival per host and thus less death^3^. The fitness value can be interpreted as a percentage reduction in death per generation. For example, a fitness of 2 would result in a 50% reduction in death rate. In addition, rather than sample selection coefficients from a distribution, we chose to have deterministic fitness values for each simulation. Therefore, neutral viruses grew at a rate *r - d* and beneficial viruses (viruses that acquired a positively selected mutation) evolved at a rate *r - d/(1+s)* dependent on the predetermined fitness value.

To generate plausible scenarios of SARS-CoV-2 evolution, we selected a set of parameters that are in line with empirical estimates. Overall, we assumed evolution can only happen within the host and that only approximately one mutation, if any, is likely to be transmitted per infected individual. Therefore, we chose *r, d*, and fitness values to be within reasonable approximations to the basic reproduction number (*R0*) or effective reproduction number (*Re*) of SARS-CoV-2 from March until December 2020. Under this assumption, we set *r* to 2.02 (1 + 1.02) and *d* to 1 such that fitnesses of 1.1, 1.2, 1.3, 1.4, 1.5, 1.6, 1.7, 1.8, 1.9, and 2, led to overall exponential growth rates (N_t+1_/N_t_) of approximately 0.11 (1.11), 0.19 (1.19), 0.25 (1.25), 0.31 (1.31), 0.35 (1.35), 0.40 (1.40), 0.43 (1.43), 0.46 (1.46), 0.49 (1.49), and 0.52 (1.52) in viruses with beneficial mutations. These values are consistent with moderate ranges of *R0/Re* observed during the pandemic from March to December 2020 (http://metrics.covid19-analysis.org/ or https://epiforecasts.io/). With that said, both neutral populations and populations subject to positive selection grew under exponential growth, therefore the relative difference in growth is the main component in the observed variation between neutral and positively selected subpopulations. For the mutation rate, we scaled our per base per generation mutation rate to match the number of mutations in simulated haplotype alignments to empirical aligned haplotype alignments (Figure 1f). All parameter combinations were simulated with per base per generation mutation rates of 1 × 10^−6^ and 5 × 10^−6^ for training CNNs on small genomic windows and 1 × 10^−6^, 2.5 × 10^−6^, and 5 × 10^−6^ for training RNNs on full 29903 base pair simulated genomes. We evolved populations until a population size of approximately one hundred thousand was reached.

Following each simulation, we randomly sampled a specified number of simulated viruses from the population to mimic the random sampling and testing process in real patient populations. Using a randomly generated reference genome, we then aligned the simulated viral genomic windows or complete genomes together and converted them to a binary encoded format, where 0 indicated the same base as the reference and 1 indicated a different base than the reference. Under this definition, the VAF for a given site was based on the sum of all variants different from the reference divided by the sample size. ImHapE operates at the nucleotide-level and is therefore agnostic in relation to the genetic code (for example, missense or silent mutations), which is relevant for RNA viruses where nucleotide changes can alter RNA binding and structure^19,20^.

We initially evaluated the separability of positive and neutral evolutionary simulations using site-frequency based summary statistics Tajima’s *D* (TajD) and Fay and Wu’s *H* (FWuH) (Supplementary Figure 1). By simulating 2000 viruses per parameter combination, we found that neutral evolution and positive selection were differentiable using TajD and FWuH across a range of genomic window sizes, mutation rates, and probabilities of mutations being positively selected.

We note that, although we integrate a custom simulation framework into our pipeline, any simulation output can be trained using the CNN if the sequence alignments are converted to binary-encoded NumPy files. All simulations parameters are provided in Supplementary Table 1.

### Convolutional and recurrent neural networks for evolutionary inference using aligned haplotypes

To make evolutionary inferences in phased genomes, such as SARS-CoV-2, we chose to take an image recognition approach. Previous work has shown the effectiveness of treating evolutionary inference as an image recognition problem^7,9^. To identify loci subject to positive selection in a non-recombining system, we built a tool combining evolutionary simulations, CNNs, and RNNs to perform sliding window analysis across a genome. Our method involves four main steps, (i) simulating evolution in genomic windows of length *N*, (ii) training a CNN on aligned images, genomic windows of length *N*, (iii) simulating complete 29903 bp genomes and performing sliding window analysis using trained CNN and other population genetic statistics such as Tajima’s D (TajD) and Fay and Wu’s H (FWuH), and (iv) training an RNN to predict if positive selection is acting on the non-recombining genome and the fitness (1 + s) within the population at a given time point.

To build training data for evolutionary inference in SARS-CoV-2, we simulated viral evolution in genomic windows of length *N* and in full length genomes of 29903 bp. Genomic windows were simulated to train CNNs on image-based alignments and full length genomes were simulated to train RNNs following sliding window analysis with the CNN. Genomic windows were simulated across multiple sizes (1000, 2500, and 5000 bp) and with various sorting methods (no sorting, row sorting, column sorting, row and column sorting). Sorting methods re-order the genomic windows based on the positions with the largest number of mutations (columns) and/or the based on the samples with largest number of mutations (rows). Sorting the image-based alignments based on mutation frequency has previously been shown to impact inferences in recombinant organisms^7^. For each genomic window simulation, we sampled 200 viruses from a population of one hundred thousand and then generated binary encoded images with 200 samples (rows) and 1000, 2500, or 5000 bp depending on the genomic window (columns) size. We then pooled 8000 positive selection simulations into one group and 8000 neutral simulations in another group and trained a binary CNN classifier, implemented using *Keras* in *tensorflow* v2.3.0, on the aligned haplotypes to generate a classification score, or probability estimate, for positive selection. We used a CNN architecture similar to a previous study^7^. The exact architecture of our CNN can be found in the ImHapE github repository. We evaluated each model on an independent validation set of simulated genomics windows. We found 2500 bp windows with sample (row) sorting to provide a balance between performance, with the population genetic statistics showing separability of neutral vs positive classes (Supplementary Figure 1), and retention of positional information, and chose this window size for future analysis.

As our goal was to infer signatures of positive selection within non-recombining genomes, we attempted to account for fully linked haplotype variants by first performing sliding window analyses with our CNN across simulated viral genomes and then adjusting these estimates using methods that take into account sequential or positional information, such as an RNN. To implement this, we performed sliding window analysis with our CNN across approximately 5000 simulated 29903 bp genomes. Using the mean estimates, from 10 subsamples of 1000 viruses, at each sliding window, we trained a bidirectional LSTM to predict if the genome was evolving neutrally or under positive selection, returning a probability of positive selection between 0 and 1. In addition, we evaluated the performance of bidirectional LSTMs combining CNN estimates with site frequency-based population genetic statistics such as TajD and FWuH. We evaluated all estimator combinations (e.g. [CNN, TajD, FWuH], [CNN, TajD], …) across a range of L1/L2 regularization values (0, 0.001, 0.002, 0.003, 0.005, 0.01) and selected the best performing combination or single estimator for empirical analysis. In addition, for quantifying fitness, we combined all non-overfitting models into one ensemble to average our estimates, resulting in the lowest mean absolute error (Supplementary Figure 3). To evaluate the effectiveness of our LSTM approach for both classifying positive selection and quantifying fitness, we evaluated the performance of an independent holdout set of approximately 3000 positively selected viral populations and 3000 neutrally evolving viral populations.

### Evolutionary inference in empirical SARS-CoV-2 populations

Following training and validation of our combined CNN and RNN approach, we applied our model to empirical SARS-CoV-2 data from March to July (GISAID^1^; https://www.gisaid.org/) and from April to December (COG UK^14^; https://www.cogconsortium.uk/). We collected 49,322 SARS-CoV-2 genome sequences from GISAID on August 11th, 2020. The majority of sequences corresponded to data obtained from December 2019 to July 2020. To ensure we were making accurate inferences, we employed a conservative filtering step removing any genomes that appeared to outliers based on the proportion of the genome masked (Supplementary Figure 4). For GISAID data, this corresponded to removing any sample with greater than 0.1% of the genome masked. We retained 28,350 samples in our final analysis. We then aligned each sequence to the Wuhan reference genome (NCBI RefSeq: NC_045512) using a Needleman-Wunsch rapid global alignment implemented in EMBOSS stretcher (default settings). We converted each aligned sequence to a binary encoded sequence. All sequenced sites identical to the Wuhan reference, previously marked as caution (https://github.com/W-L/ProblematicSites_SARS-CoV2), or containing a gap except for positions between 21765 - 21770 (known deletion), were labeled 0 and all sequenced sites with base calls different from the reference were labeled 1.

Because our CNN was trained on alignments of 200 samples or sequences, we required approximately 200 samples per geographic region for analysis. In addition, as we were looking to analyze positive selection estimates over time, we required 200 samples per time point. Following filtering of the data, the GISAID regions of North America, Europe, and Asia had a sufficient (>= 200) number of samples per month between March to July 2020. We then performed sliding window analysis, with a step size of 50 bp, across both the North America, Europe, and Asia SARS-CoV-2 samples using our combined CNN and LSTM approach. To generate confidence intervals, we subsampled the population 25 times at each sliding window using the mean, 2.5%, and 97.5% values for our positive selection and fitness estimates.

In addition to GISAID samples, we collected 112,212 aligned viral genomes from COG UK on December 21st, 2020. We only kept samples that were available across both GISAID and COG UK data sources at the time of this study. Similar to GISAID, we applied a conservative filtering approach removing any genomes that appear to be outliers based on the proportion of the genome masked. For COG data, this corresponded to removing any sample with greater than 1.8% of the genome masked. We retained 66,495 samples in our final analysis. As we were interested in studying potential positive selection beyond the time points investigated in the GISAID samples, we selected an England reference genome to call variants using a virus that harboured the known fixed D614G haplotype. The reference ID used to call variants in the COG samples was England/LOND-D51C5/2020 from week 16. Given the effective sequencing program by COG UK we were able to implement our CNN and LSTM approach at a weekly time scale resolution from approximately week 16 to week 50 of 2020. We removed estimates from any time point with less than 200 samples as we required a minimum of 200 samples in our alignments.

To ensure our estimates weren’t biased by fixed variants (VAF = 1), we excluded fixed sites from our analyses. Technically, fixed variants provide no relative fitness advantage as they are present in all individuals within a population but have the potential to upwardly bias fitness estimates as positive selection increases probability of fixation relative to neutral evolution. To remove fixed variants in the alignments, we converted the fixed site in all samples at a given time point to 0 when performing sliding window analysis.

### General statistical and data analyses

Simulation framework was developed using *python* v3.6.0. All data processing, visualization, and analysis was performed using *python* v3.6.0 or *R* v4.0.3. All machine learning models were implemented in *Keras* using *tensorflow* v2.3.0 or *scikit-learn* v0.24.0. TajD and FWuH were computed using a custom python function. All code used in this study is available on github (see Data/Code availability).

